# Molecular dynamics simulations and functional studies reveal that hBD-2 binds SARS-CoV-2 spike RBD and blocks viral entry into ACE2 expressing cells

**DOI:** 10.1101/2021.01.07.425621

**Authors:** Liqun Zhang, Santosh K. Ghosh, Shrikanth C. Basavarajappa, Jeannine Muller-Greven, Jackson Penfield, Ann Brewer, Parameswaran Ramakrishnan, Matthias Buck, Aaron Weinberg

## Abstract

New approaches to complement vaccination are needed to combat the spread of SARS-CoV-2 and stop COVID-19 related deaths and long-term medical complications. Human beta defensin 2 (hBD-2) is a naturally occurring epithelial cell derived host defense peptide that has antiviral properties. Our comprehensive *in-silico* studies demonstrate that hBD-2 binds the site on the CoV-2-RBD that docks with the ACE2 receptor. Biophysical and biochemical assays confirm that hBD-2 indeed binds to the CoV-2-receptor binding domain (RBD) (K_D_ ∼ 300 nM), preventing it from binding to ACE2 expressing cells. Importantly, hBD-2 shows specificity by blocking CoV-2/spike pseudoviral infection, but not VSV-G mediated infection, of ACE2 expressing human cells with an IC_50_ of 2.4± 0.1 μM. These promising findings offer opportunities to develop hBD-2 and/or its derivatives and mimetics to safely and effectively use as novel agents to prevent SARS-CoV-2 infection.

## INTRODUCTION

The ongoing COVID-19 pandemic, the result of infection by SARS-Coronavirus-2 (CoV-2), continues to infect people worldwide; having claimed over 1.75 million lives (Johns’ Hopkins University) as of late December 2020. While the first vaccines are now being administered, albeit initially to a select population, the virus continues to evolve in significant ways. This situation requires the discovery of novel therapeutic approaches, possibly to be used independently or in conjunction with existing approved regimens, to impede the virus’ relentless spread.

All coronaviruses, including CoV-2, express the all-important S (Spike) protein that gives these viruses the characteristic corona or crown appearance (Siu et al., 2008; Yoshimoto, 2020). The S protein is responsible for binding to the host cell receptor followed by fusion of the viral and cellular membranes, (Walls et al., 2016). To engage a host cell receptor, the receptor-binding domain (RBD) of the S protein undergoes hinge-like conformational movements that transiently hide or expose its determinants for receptor binding (Wrapp et al., 2020). Structural fluctuations of the RBD, relative to the entire S protein, enable exposure of the receptor-binding motif (RBM), which mediates interaction with the receptor angiotensin-converting enzyme 2 (ACE2) on the host cell (Lan et al., 2020; McCallum et al., 2020; Walls et al., 2020; Yan et al., 2020). Since this is believed to be the critical initial event in the infection cascade, the RBD has been proposed as a potential target for therapeutic strategies (Tai et al., 2020). The high degree of dynamics of the RBD:ACE2 complex (Brielle et al., 2020; Ghorbani et al., 2020; Spinello et al., 2020; Xiong et al., 2020), suggests that binding of small flexible proteins and peptides may inhibit Spike protein:host cell receptor interactions, which can be interrogated by computational modeling and simulations most suitable for exploring these interactions (Amaro and Mulholland, 2020).

Nature’s own antimicrobial peptides (AMPs) have been proposed as multifunctional defenses that participate in the elimination of pathogenic microorganisms, including bacteria, fungi, and viruses (Diamond et al., 2009). Exhibiting antimicrobial and immunomodulatory properties, AMPs have been intensively studied as alternatives and/or adjuncts to antibiotics in bacterial infections and have also gained substantial attention as anti-viral agents (Mulder et al., 2013). Human beta defensins (hBDs), the major AMP group expressed naturally in mucosal epithelium, provide a first-line of defense against various infectious pathogens, including enveloped viruses (Leikina et al., 2005; Quiñones-Mateu et al., 2003; Ryan et al., 2011).

The hBDs are cationic peptides, which assume small β-sheet structures varying in length from 33 to 47 amino acid residues and which are primarily expressed by epithelial cells (Bensch et al., 1995; Harder et al., 2001; Harder et al., 1997; Schibli et al., 2002). HBD-2 has been shown to express throughout the respiratory epithelium from the oral cavity to the lungs and, it is believed that this defensin plays a very important role in defense against respiratory infections (Diamond et al., 2008). Altered hBD-2 expression in the respiratory epithelium is known to be associated with the pathogenesis of several respiratory diseases such as asthma, pulmonary fibrosis, pneumonia, tuberculosis and rhinitis, (Diamond et al., 2008; Doss et al., 2010; Ooi et al., 2015; Rivas-Santiago et al., 2005; Semple and Dorin, 2012). HBD-2 has been demonstrated to inhibit human respiratory syncytial virus (RSV) infection by blocking viral entry through destabilization/disintegration of the viral envelope (Kota et al., 2008). It might also have important immunomodulatory roles during coronavirus infection as well, as hBD-2 conjugated to the MERS receptor binding domain (RBD) has been reported in a mouse model to promote better protective antibodies to RBD than RBD alone (Kim et al., 2018).

In the present study, we examined the ability of hBD-2 to act as a blocking agent against CoV-2. HBD-2 is an amphipathic, beta-sheeted, highly cationic (+6 charge) molecule of 41 amino acids, and is stabilized by three intramolecular disulfide bonds that protects it from degradation by proteases (Sawai et al., 2001). The protein has been studied before with molecular dynamics simulations, (Yeasmin et al., 2018) (Barros et al., 2020; Ghorbani et al., 2020; Spinello et al., 2020). Through extensive *in silico* docking and molecular dynamic simulation analyses we report herein that hBD-2 binds to the receptor binding motif (RBM) of the RBD of CoV-2 that associates with the ACE2 receptor. Biophysical and biochemical studies confirmed that hBD-2 binds the RBD and also prevents it from binding ACE2. Moreover, by utilizing a physiologically relevant platform, we revealed that hBD-2 effectively blocks CoV-2 spike expressing pseudovirions from entering ACE2 expressing human cells. Harnessing the utility of naturally occurring AMPs, such as hBD-2, and their derived smaller peptides, could be a viable approach at developing novel CoV-2 therapeutics.

## RESULTS

### Interrogating the interaction of SARS-CoV-2 RBD with ACE2 and hBD-2 using *in silico* docking and molecular dynamics simulations

#### RBD:ACE2 complex

We began our *in silico* work by running, as a reference, a 50 ns all-atom molecular dynamics (MD) simulation of the ACE2:RBD complex. The final structure was compared with the initial experimental crystal structure (Lan et al., 2020), as shown in Figure S1A. Only small deviations are seen in some of the loop regions and at the N- and C-termini of both proteins; the overall rms deviation (RMSD), of the structure, calculated for backbone Ca atoms, is around 1.2 Å, for ACE2, around 2.1 Å for the RBD and around 2.4 Å for the complex (Figure S1B) [Supplementary information]. The result of calculating the rms fluctuation (RMSF) for the Ca atom of each residue in the RBD and in ACE2 is shown in Figure 1A (Left and Right). Overall, the main-chain fluctuations in the RBD and ACE2 are small with a magnitude of around 0.6 Å for the most structured, α-helical and β-sheet parts. As can be seen, the loop regions are more flexible, having a higher RMSF of up to 4 Å. The difference in fluctuations between ACE2 and RBD in their bound and free states in solvent are shown in Figure 1B. As is usually expected, most regions at the RBD:ACE2 interaction interface become less flexible (shaded in blue), while other changes, including increases in fluctuations (shaded in red) are seen further away from the interface, consistent with the recent description of allostery in the spike protein (Gross et al., 2020; Ray et al., 2020). Upon complex formation, the RBD and ACE2 proteins form intermolecular hydrogen bonds, which is one of the driving forces for their binding. These bonds, calculated over the course of the 50 ns simulation, are plotted in Figure 1C. In the first 15 ns, the average number of hydrogen bonds fluctuates between 2-7, but settles at a slightly lower number, 2-5, at the end. Importantly, these bonds are highly dynamic with occupancy between 20-40%. Hydrogen bonds with good persistency are listed in the table in Figure 1. ACE2 residues Lys353 and Gly502, Tyr83 and Asn487, Asp30 and Lys417 formed hydrogen bonds with duration of at least 34%. In total, 7 of 9 H-bonds of the RBD:ACE2 interface in the crystal structure (Lan et al., 2020) are populated with reasonable occupancy in the simulations. Similar behavior has been seen in other simulations (Ghorbani et al., 2020; Spinello et al., 2020) with the difference likely explained by solution vs. crystallization conditions. Water molecules were observed at the interface in other simulations and are likely bridging the interactions (Malik et al., 2020), also underscoring the dynamic nature of the interactions (see below). To further indicate the overall stability of the interface in the simulations, we calculated the solvent accessible surface area, which is buried between the RBD and ACE2 proteins in the complex. During 50 ns, this buried surface area fluctuates between a minimum of 750 Å^2^ to a maximum of 1000 Å^2^, but this is maintained at an average of ∼900 Å^2^ over the last 25 ns of the trajectory. We also calculated the distance map between ACE2 and RBD atoms, which are closer than 5 Å on average as a reference (see below).

**Figure 1.**
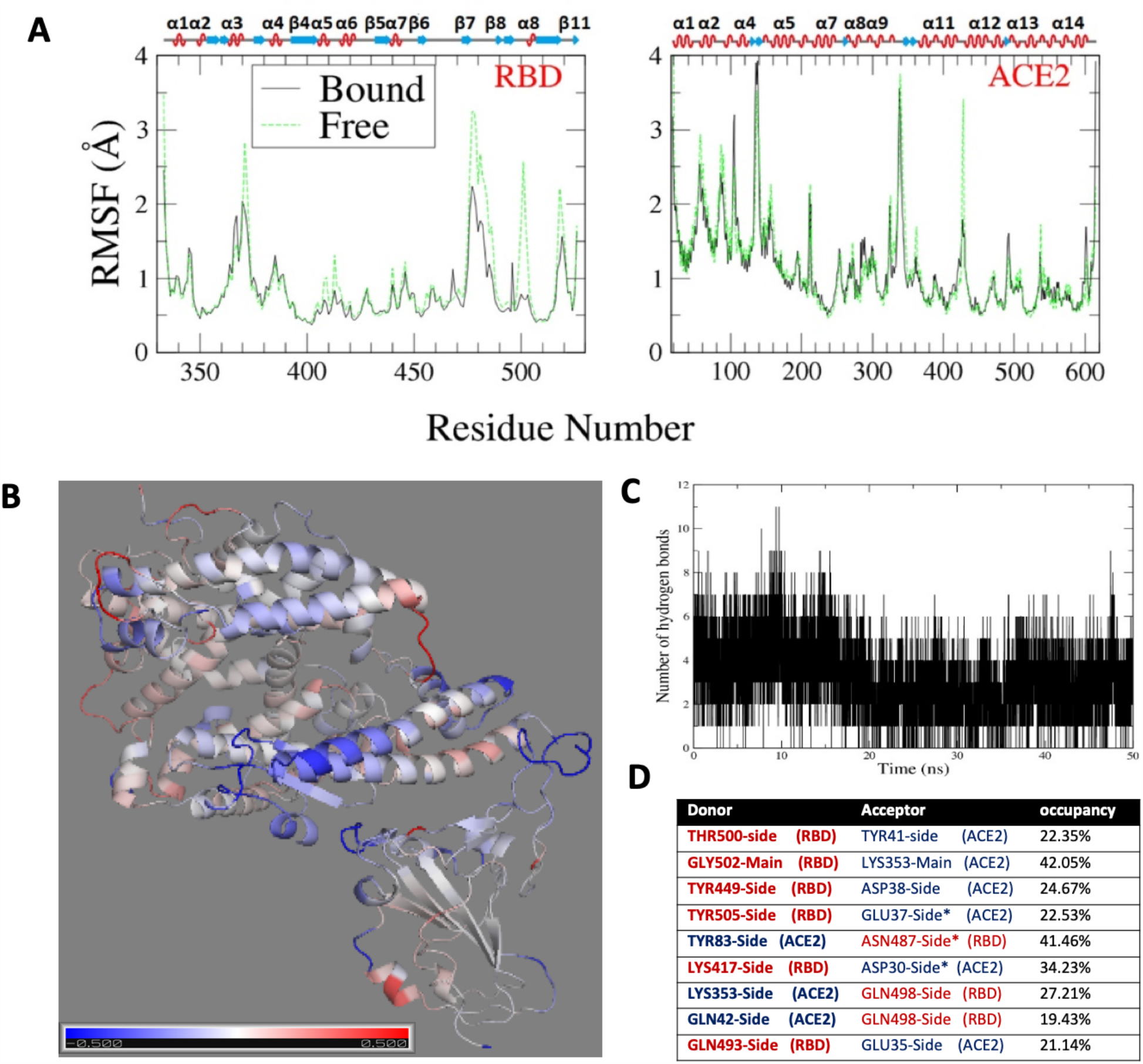
Molecular dynamics simulations of RBD:ACE2 (as a reference) show protein complex is stable. (A) RMSF of RBD (left) and ACE2 (right) in the complex over 50 ns in comparison with values for the unbound (free) proteins; the secondary structure of ACE2 and RBD are indicated. (B) Difference in RMSF between bound and free proteins. The data are mapped to the cartoon representation of the complex with color bar (Bottom) indicating the range of −0.5 Å (in blue) to 0.5 Å (in red) (C) Number of hydrogen bonds for the RBD bound to ACE2 over the course of the simulation. (D) Table of most prominent h-bonds and their occupancy

#### RBD:hBD-2 (monomer)

In order to explore the initial possible bound structures between the two proteins, we carried out docking with Cluspro and Haddock (see Methods in supplement). The best predicted models were used as starting structures for all-atom MD, as above; however, since the initial docked structures are not well converged, we carried out the simulations for up to 500 ns. We also ran repeat simulations with different starting seeds (initial velocity assignments). The simulations performed are summarized in Table S1 [Supplementary information]. In Figure 2A we present the most converged and apparently stable trajectory, showing the initial structure when compared to the last structure (after 500 ns). Slight rotation of hBD-2 relative to the initial structure is indicated at 75 ns by the transition of RMSD when plotted as a function of simulation time (Figure 2B); however, for the remaining 427 ns, hBD-2 stayed in the same position. Analysis of the other three trajectories is provided in Figure S2.

**Figure 2.**
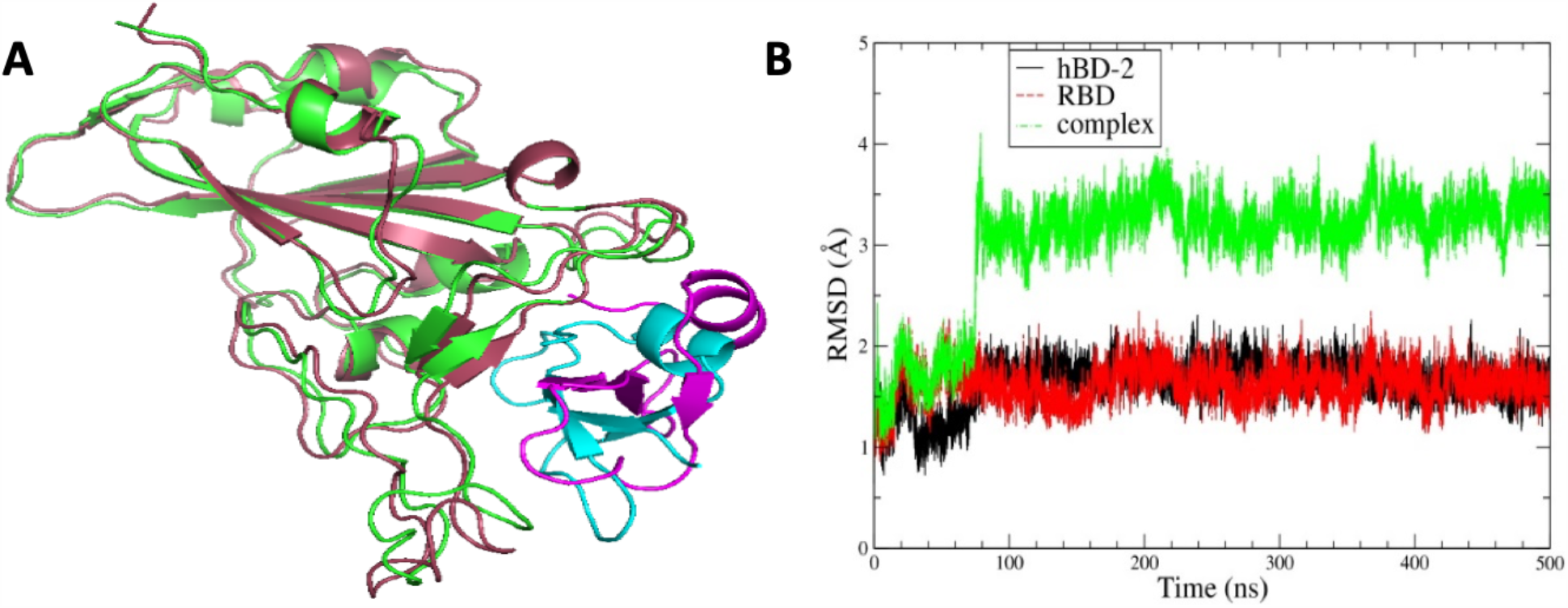
Cartoon representation of RBD:hBD-2. (A) Comparison of the initial and last structure after 500 ns simulation (shown in cyan for hBD-2 and green for RBD and shown in magenta for hBD-2 and raspberry for RBD respectively) after 500 ns all-atom MD simulations for the RBD:hBD2 complex (B) RMSD of proteins in the complex and of the complex itself.

The comparison of main-chain fluctuations in RBD and hBD-2, between their bound and free states is shown in Figure 3A. Overall, the binding region becomes less flexible on the RBD in similar key loop regions whose dynamics are dampened by ACE2 binding, while on the side of hBD-2 a significant number of main-chain sites also see their fluctuations decreased. The results are mapped to the final structure of the trajectory in Figure 3B. As above, we calculated the number of intermolecular hydrogen bonds formed between hBD-2 and RBD over the course of the trajectory (Figure 3C). They are fewer, with an average 4 ± 1, compared to those bridging the RBD:ACE2 complex. Similarly, with the exception of the hBD-2 residue Arg23, which forms a hydrogen bond with the ACE2 residue Glu484 greater than 50% of the time, the occupancy of other hydrogen bonds is reduced compared to the reference complex. As before, the occupancy of these interactions is not 100%; i.e., more like 30%, suggesting that they are somewhat dynamic (see discussion below) and are accompanied by indirect H-bond interactions with water molecules near or at the interface bridging the interactions (Malik et al., 2020). Both of these features were also found in simulations of the RBD:ACE2 interaction, as already noted; however, the dynamics of these interactions appear to be more prevalent in the RBD:hBD-2 interaction. As might be expected for the cationic hBD-2, the positively charged sidechains are a prominent feature in the interactions, especially Arg22 and Arg23. The RBD residues most persistently involved in the interaction with hBD-2 are shown in the table of Figure 3. With the exception of Gln498, the interaction between hBD-2 and RBD involves amino acids that are within a few residues of those that are involved between ACE2 and RBD and cover a good proportion of the same interface area.

**Figure 3.**
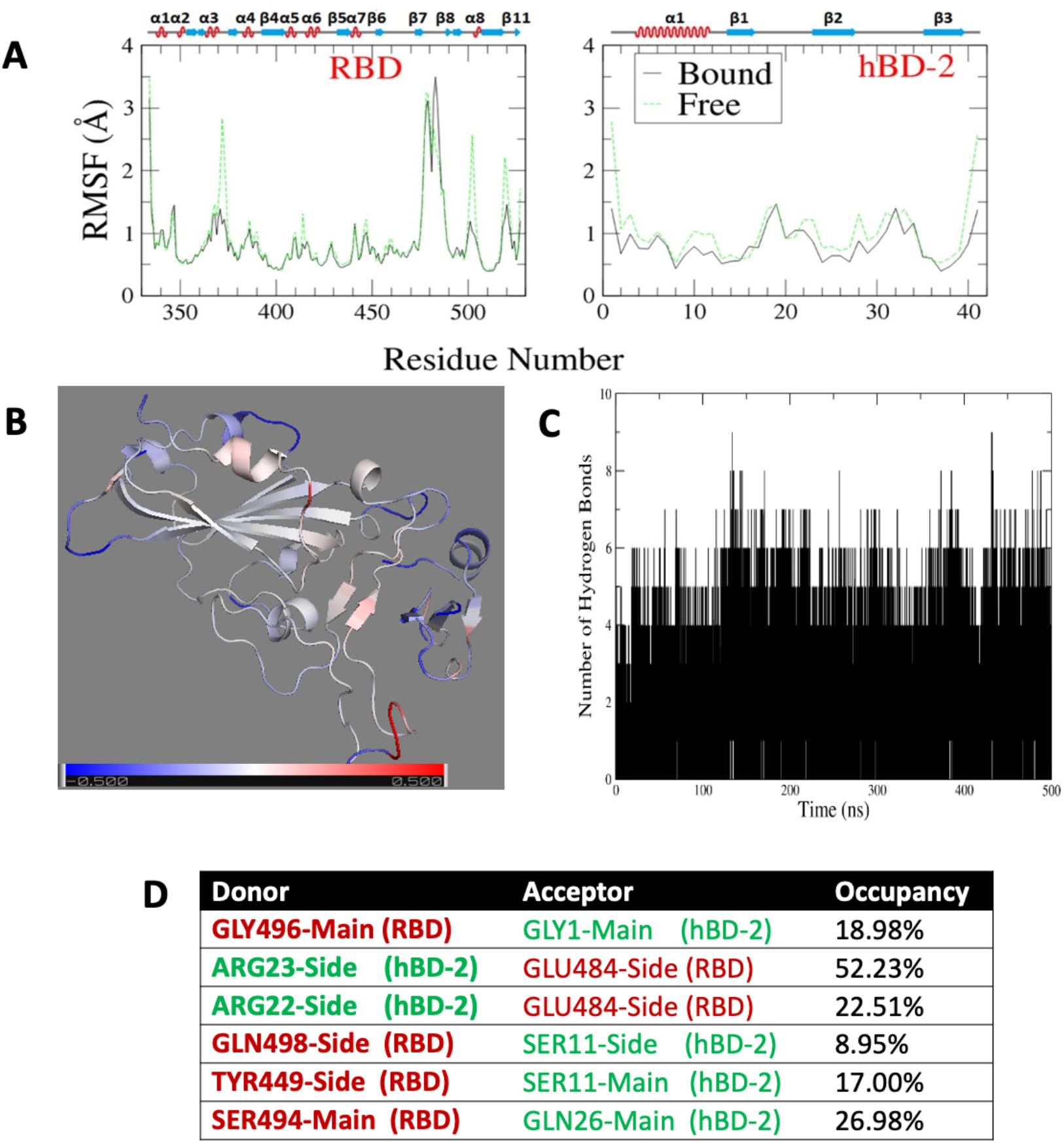
The RBD and hBD-2 proteins retain considerable dynamics as a complex. (A) RMSF of RBD (left) and hBD-2 (right) in the complex over 500 ns in comparison with values for the unbound (free) proteins; the secondary structure of ACE2 and RBD are indicated (B) Difference in RMSF between bound and free proteins. The data are mapped to the cartoon representation of the complex with color bar (Bottom) indicating the range of −0.5 Å (in blue) to 0.5 Å (in red) (C) Number of hydrogen bonds for the RBD bound to hBD-2 over the simulation. (D) Table of most prominent h-bonds and their occupancy.

The persistency of the complex is also confirmed in the changes in accessible surface area, which is buried between the two proteins, and fluctuates moderately around a value of 700 ± 150 Å^2^. The value is smaller than that of ACE2 (900 Å^2^), indicating that less area is covered. This is expected since the hBD-2 protein is considerably smaller than the RBD. A distance map, comparing residues which are on average closer than 5 Å in the RBD:ACE2 and RBD:hBD-2 complexes is shown in Figure 4. For the RBD:ACE2 interaction (Figure 4A), residues 20 to 45, 75 to 85, as well as a short stretch of residues around 327, 355 and 387 on ACE2 bind with the RBD, whose binding interface ranges from residue 445 to 505. Some of the RBD residues are in loop regions; e.g., 404 and 417, which also come close to ACE2 over the course of the 50 ns simulation. The contact analysis for the RBD:hBD-2 complex over the course of the 500 ns simulation is shown in Figure 4B. Remarkably, in comparison with the RBD:ACE2 complex, essentially all residues of the RBD which contact ACE2, either the same ones or their close neighbors, are also in contact with hBD-2. However, there are some subtle shifts. For example, RBD residues 475-478 make contact with ACE2 but not with hBD-2, where these interactions may have shifted to residue 473. Also, a regional area of residues 438-444 contacts hBD-2, which is not seen with ACE2. These contacts may be absent in the RBD:ACE2 complex because it is less dynamic, and only sampled for 50ns. Alternatively, they may provide a mechanistic entry for hBD-2 in replacing/competing away ACE2 from the spike trimer.

**Figure 4.**
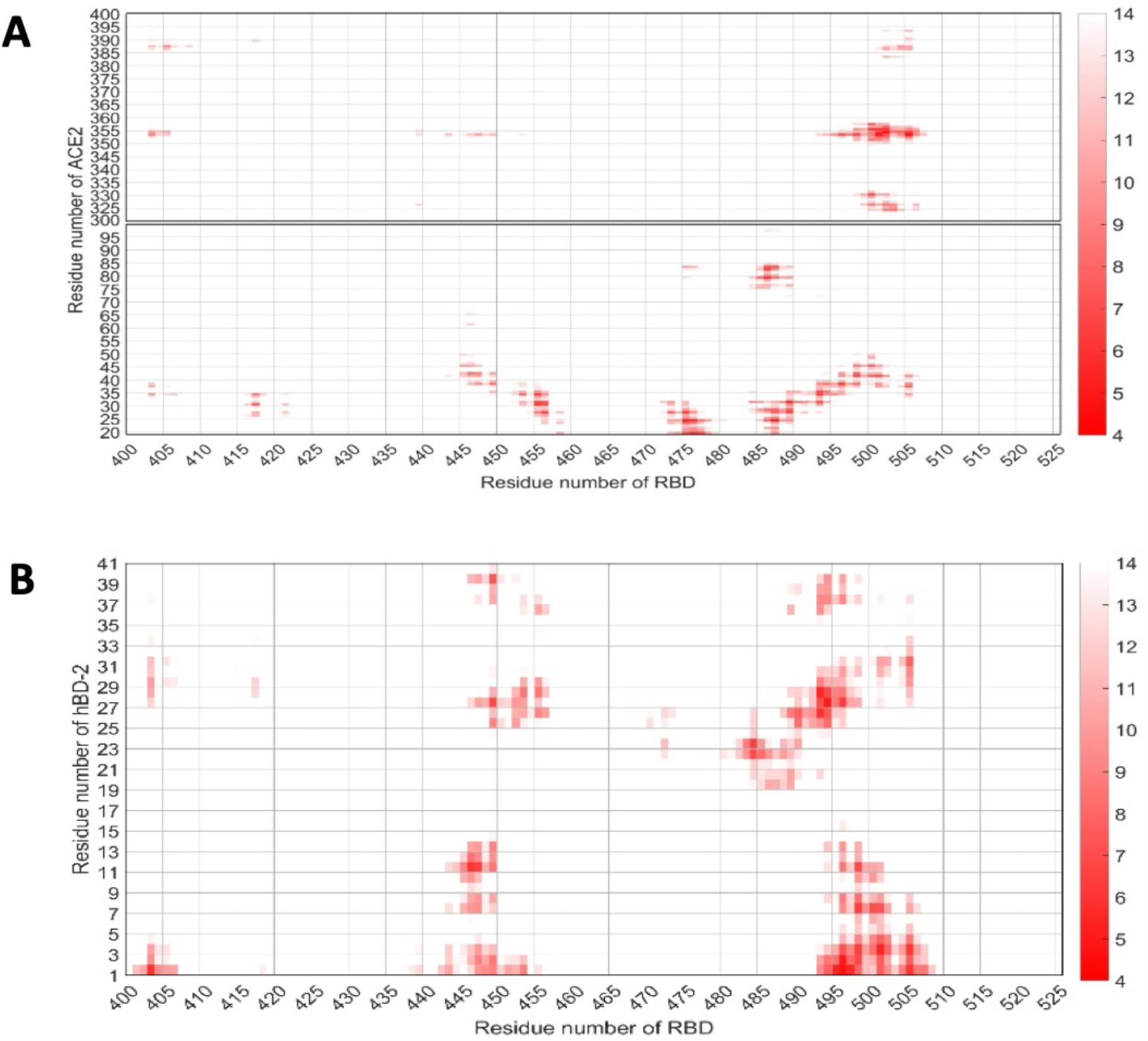
Similar regions/residues are involved in RBD contact with ACE2 as with hBD-2. (A) Distance map of inter-protein contacts in (A) the RBD:ACE2 complex and (B) the RBD:hBD-2 complex with distances color coded by average proximity over the length of the simulations (see color scale, right).

In order to confirm consistent binding of hBD-2 to RBD, we started simulations from the same initial structure of Figure 2 and repeated the simulation for three more times, each with a different random seeds (simulation details are shown in Table S1, and results are shown in Figure S2). These simulations are consistent with the results above in terms of the RMSD and the average surface area buried. The change in fluctuations in forming the complex varied. The average numbers of hydrogen bonds, around 2 ± 1 at any one time, are slightly less; however, as stated above, Arg22 and Arg23 are the major residues on hBD-2 contributing to the formation of hydrogen bonds.

#### RBD:hBD-2 (dimer)

Although the affinity of hBD-2 for dimerization is modest (Hoover et al., 2000), it is possible that binding to the RBD stabilizes the dimeric form. We, therefore, also docked the hBD-2 dimer to the RBD and carried out simulations. The initial and final structure comparison of one of the simulations is shown in Figure S3 A and the RMSD is plotted as a function of simulation time (Figure S3 B). The comparison of the RMSF of the hBD-2 dimer and RBD in their bound state with their fluctuations in their free state is shown as well (Figure S3). Intriguingly, when compared to hBD-2 monomer binding, a few regions of the RBD do not diminish as much in flexibility, while some actually become more flexible. The buried surface accessible area is slightly larger (about 10% larger) for the dimer compared to monomer binding, confirming that interactions to both units of the dimer from the RBD exist. The distance map is given in Figure S4. As shown, both units of the hBD-2 dimer can bind with the RBD in the residue range of 445 to 500. Mostly dimer associated residues from 17 to 24 are in tight and close contact with the RBD. The number of hydrogen bonds formed between the hBD-2 dimer and the RBD are similar to those formed by the monomer and the RBD (Figure 3C and Figure S4A). Again the hBD-2 Arg23 is the most prominent interacting residue. In fact, unit 1 of the dimer can form more hydrogen bonds with the RBD, and also one hydrogen bond from unit 2 is prominent, again involving its Arg23, this time to Glu406 on RBD, which is outside the region typically interacting with ACE2. Remarkably, the persistency of the hydrogen bonds is increased from ∼ 30% in the monomer to ∼50% in the dimer (shown in Figure S5), suggesting overall that binding of a dimeric hBD-2 may be favorable.

### RBD:hBD-2-interaction energy calculation

Due to the caveats associated with calculations of free energy estimations from trajectories such as the ones run for this study, we carried out the binding interaction energy calculation for RBD binding with ACE2 and hBD-2 monomer/dimer, respectively, using the popular GBSA method (see Materials & Methods section). We report the average energies and standard deviations as a histogram in Figure S6. These interaction energies have similar values and all are slightly negative. Comparing the binding energy between RBD with ACE2 and with hBD-2 monomer/dimer, the average binding energy of the RBD with ACE2 is −37 ± 8 kcal/mol whereas average binding energy of RBD with hBD-2 dimer is −34 ± 8 kcal/mol, and similarly for RBD binding with hBD-2 monomer. However, it is likely that the entropy change upon binding RBD is significantly more favorable for binding to hBD-2 than binding to ACE2 since the former is more dynamic in the bound state, giving less of an entropy penalty upon binding. In fact this latter indication suggests that peptides, which are initially unstructured in the unbound state could also maintain considerable flexibility in the bound state and may thus be powerful antagonists of the RBD:ACE2 interaction. Detailed thermodynamics analyses, both experimental and computational are needed to clarify this point. Irrespective of these estimated numerical values, the calculations suggest that hBD-2 at a sufficiently high concentration should be able to block the binding of RBD with ACE2. Our experimental analysis with RBD:hBD-2 interactions using purified proteins and the spike-pseudovirion assay suggests such a concentration is likely to be in the vicinity of the IC_50_ of 2.4 µM.

### Experimental studies confirming the binding of hBD-2 with the RBD

We used multiple experimental approaches to confirm the *in silico* findings of hBD-2 and SARS-CoV-2 RBD binding. Microscale Thermophoresis (MST) showed that CoV-2 RBD interacts with recombinant hBD-2 (rhBD-2) with a dissociation constant of ∼300 nM (Figure 5A). This interaction is weaker (> 3 μM) when hBD-2 loses its natural conformation under disulfide bond reducing conditions (Figure 5A). We then followed up using a functional ELISA assay, and found that rhBD-2 bound to immobilized RBD in a linear range (over concentrations of 1.5 to 100 nM), as detected by biotinylated anti-hBD-2 detection antibodies (Figure 5B). We then examined the binding of rhBD2 and recombinant histidine tagged-RBD (His-RBD) derived from our expression system for codon optimized CoV-2 RBD (see Materials and Methods) by co-immunoprecipitation. By incubating rhBD-2 with His-RBD at a ratio of 1.5:1.0, followed by nickel bead immunoprecipitation of His-RBD and probing for hBD-2 in Western blots, we found significant binding of hBD-2 to His-RBD (Figure 5C). Control Western blots showed only modest background binding of hBD-2 in the absence of RBD, thereby confirming the specificity of the RBD:hBD-2 interaction (Figure 5C).

**Figure 5.**
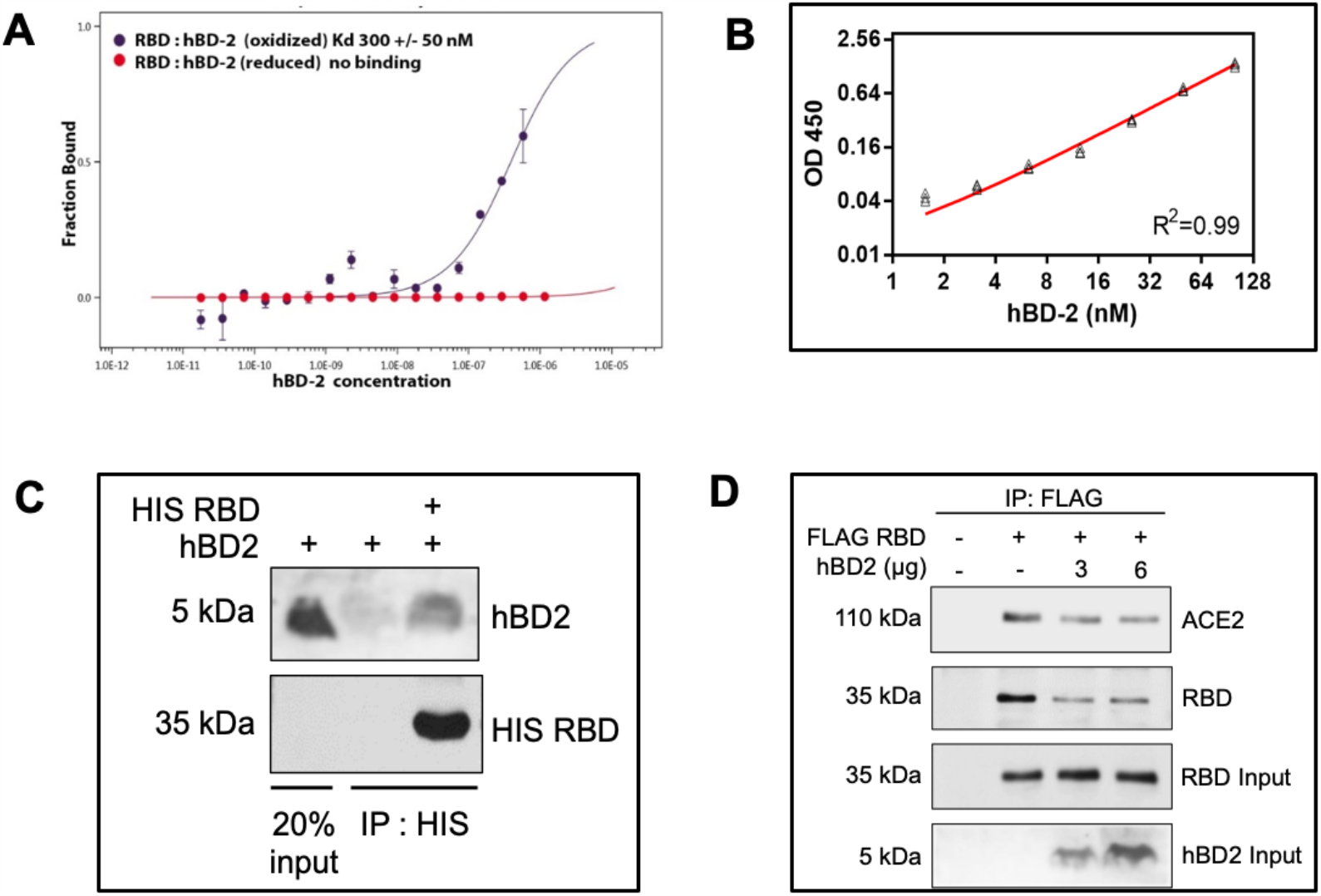
Biophysical and biological assays demonstrating hBD-2 binding to RBD. (A) Concentration dependent binding of recombinant hBD-2 (rhBD-2) to fluorescently labeled recombinant RBD (rRBD), as measured by miscroscale thermophoresis. HBD-2 was used under oxidizing (black data points) and under reducing conditions (red). (B) Functional ELISA assay showing that rhBD-2 binds to immobilized rRBD with a linear range of concentrations (1.5 to 100nM). (C) Recombinant His-RBD (5 µg) and hBD-2 (7.5 µg) were incubated as described in Methods and precipitated with Ni-NTA beads to pulldown His-tagged-RBD. Co-precipitation of hBD-2 was assessed by Western blotting. Lane 1 shows 20% input of hBD-2 and lane 2 shows Ni-NTA precipitation to examine background binding of hBD-2 to the beads. Data is representative of three independent experiments. (D) ACE2 HEK 293T cells were incubated with FLAG-RBD, with and without hBD-2 at indicated concentrations. Anti-FLAG immunoprecipitation was performed to precipitate ACE2 bound to FLAG-RBD and to assess the effect on hBD-2 addition of RBD:ACE2 binding. Data is representative of two biological replicates.

### HBD-2 blocks the binding of RBD with cellular ACE2

Next, we examined whether rhBD-2 can interfere with the binding of RBD to the host ACE2 receptor. We utilized HEK 293T cells that overexpress the human ACE2 receptor in the assays and incubated these cells with FLAG-RBD containing culture supernatant with and without rhBD-2. We immunoprecipitated RBD through the FLAG tag and examined the co-precipitation of ACE2. We found that FLAG-RBD effectively precipitated ACE2 and the addition of hBD-2 competitively decreased RBD-ACE2 binding (Figure 5D). RBD levels were also decreased in the immunoprecipitate upon rhBD-2 addition, further suggesting a direct interaction of RBD with rhBD-2, thereby preventing RBD-ACE2 binding (Figure 5D).

### HBD-2 specifically inhibits SARS-COV-2 spike-mediated pseudoviral infection

After discovering that rhBD-2 binds RBD and competitively inhibits RBD binding to ACE2, we investigated whether rhBD-2 can inhibit spike mediated pseudoviral entry into ACE2 expressing cells. A luciferase reporter expressing CoV-2 spike-dependent lentiviral system (Crawford et al., 2020) was used to study the competitive inhibitory effects of rhBD-2 on CoV-2 spike-mediated infection. We infected ACE2 expressing HEK 293T cells using the pseudotyped virus and found substantial luciferase activity in a viral dose dependent manner (Figure 6A). Next, we studied the effect of rhBD-2 on spike-dependent viral infection of ACE2/HEK 293T cells by luciferase activity and found that hBD-2 decreased the spike mediated pseudoviral infection (Figure 6B and 6C).

**Figure 6.**
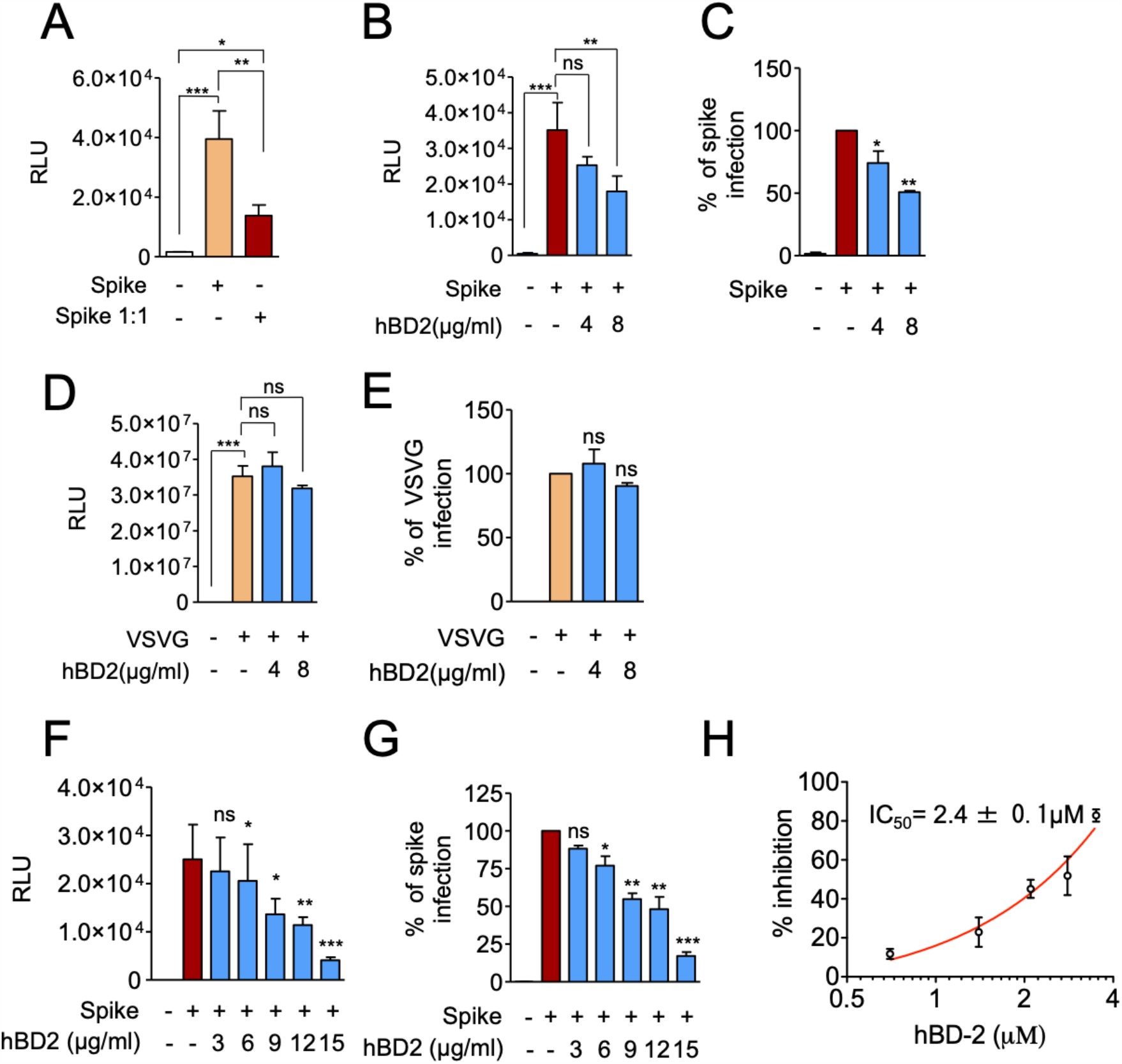
HBD2 inhibits CoV-2 spike-pseudotyped virus entry into ACE2 293T cells. (A) ACE2 HEK 293T cells were infected with CoV-2 Spike-pseudotyped virus and luciferase activity was assessed at 48 hours post infection. (B) Effect of hBD2 on CoV-2 Spike-pseudotyped virus cell entry was assessed as in A. (C) Percentage infection was calculated from the RLU values in (B) taking spike alone group as 100%. (D) Effect of hBD2 on VSVG-pseudotyped virus entry was assessed as in (A). (E) Percentage infection was calculated from the RLU values in (D) taking VSVG alone group as 100%. (F) Titration of hBD2 concentration (0-15 µg/ml) on spike-mediated pseudovirus entry and luciferase activity. (G) Percentage of spike infection was calculated from the RLU values in (F) taking spike alone group as 100%. (H) hBD2-mediated percent inhibition of spike-viral entry and IC_50_ was calculated by plotting hBD2 concentration (in µM) against % inhibition observed. Values given are Mean ± SEM of two independent experiments done in triplicates. ***p < 0.001, **p < 0.01, *p < 0.05, and ns (non-significant) against CoV-2 spike-pseudotyped virus alone treated group.

To further validate that the inhibitory effect of hBD-2 is specific to a spike-mediated infection, we used a virus pseudotyped with vesicular stomatitis virus glycoprotein (VSVG) as an independent control. Viruses pseudotyped with VSVG are pantropic; i.e., they can infect all cell types (Lever et al., 2004), and do not depend on ACE2 for entry. We obtained significant infection of ACE2/HEK 293T cells using VSVG pseudotyped virus without or with the addition of rhBD-2 (Figure 6D and 6E), thereby demonstrating the specificity of hBD-2 in blocking CoV-2 spike glycoprotein mediated infection of ACE2 expressing cells. We then inquired if increased inhibition of spike mediated pseudoviral entry was directly proportional to increased concentration of hBD-2. We discovered that indeed there was a clear hBD-2 dose response inhibition of pseudoviral entry (Figure 6F and 6G), and that the inhibitory concentration_50_ (IC_50_) was approximately 2.4 ± 0.1 µM (Figure 6H). At a concentration of 15 µg/ml rhBD-2 decreased the spike-mediated pseudoviral infection by over 80% (Figure 6H).

## DISCUSSION

The human body expresses over a hundred AMPs that are found in either intracellular granules of professional phagocytes and/or in epithelial cells of mucosa lining our external and internal surfaces (Dawgul et al., 2016). Beta defensins and LL-37, the only member of the cathelin AMPs expressed in humans, are localized to the mucosa of the oral cavity, nares and upper airway (Diamond and Ryan, 2011; Ghosh et al., 2007; Khurshid et al., 2017; Lee et al., 2002; Mathews et al., 1999; Singh et al., 1998); i.e., sites deemed vulnerable to CoV-2 entry and initial infection. Indeed, these two types of AMPs, part of the epithelial cell’s arsenal of innate responses used to defend against viral challenges at mucosal sites, have been shown to interrupt viral infection of various viruses, including coronaviruses (Kim et al., 2018). However, when a mucosal site becomes overwhelmed by a microbial threat, replenishing the AMP armamentarium locally after initial release; i.e., time from transcriptional activation, translation, post-translational modification to rerelease, takes multiple hours and makes bystander cells more vulnerable to viral infection. Moreover, if a microbial threat can inhibit production or release of these AMPs, it renders this innate defense useless. To overcome this, the AMPs or their mimetics, if administered exogenously in high enough concentrations, could be a sound therapeutic strategy to protect the host at vulnerable mucosal sites without eliciting an unwanted immunological response against the agent. Interestingly, these same AMPs have been shown to be released by human mesenchymal stem cells (hMSCs) (Krasnodembskaya et al., 2010; Sutton et al., 2016), recently repurposed to treat COVID-19 patients. While hMSCs have been shown to contribute to the recovery of severely ill CoV-2 infected patients (Moll et al., 2020; Tsuchiya et al., 2020), the role that AMPs play and the mechanism by which hMSCs ameliorate symptoms of COVID-19 remains to be determined. However, the modulation of severe inflammation and microbicidal activity related to pulmonary disease are outcomes attributable to these AMPs (Alcayaga-Miranda et al., 2017; Chow et al., 2020; Krasnodembskaya et al., 2010; Sutton et al., 2016).

We chose to interrogate hBD-2 for its ability to block CoV-2 from infecting vulnerable cells because of its innate role in protecting the oral cavity and the upper airway, and because its mouse ortholog has been shown to inhibit other coronaviruses (Zhao et al., 2016). The computer simulations that we ran of hBD-2 and the RBD showed remarkable stability of the complex even after 500 ns. There was also a clear overlap of binding sites when compared to the RBD:ACE2 complex, as verified by analysis of protein-protein residue contact distance maps. Multiple methods involving MST, ELISA and immunoprecipitation followed by western blotting independently verified that hBD-2 binds to the RBD, thereby validating our *in silico* data. Competitive inhibition assays were able to show that hBD-2 reduced RBD:ACE2 binding by removing RBD from solution, which would otherwise be available for binding ACE2. Finally, by incorporating a luciferase reporter expressing CoV-2 spike-dependent lentiviral system (Lever et al., 2004), we demonstrated that hBD-2 inhibited viral entry into ACE2 expressing HEK 293T cells in a dose dependent manner, with an IC_50_ of ∼2.4 µM. This concentration is much less than most other inhibitory concentrations attributed to hBD-2 antimicrobial activity (Joly et al., 2004) and points to a favorable affinity, and possibly also avidity, of the interaction between hBD-2 and the RBD. Interestingly, hBD-2 begins to show hemolytic activity at a concentration 30 times greater (70 µM) than our IC_50_ (Koeninger et al., 2020), and shows no signs of cytotoxic effects against various other human cells (Warnke et al., 2013) at over twice our IC_50_ (Herrera et al., 2016; Mi et al., 2018; Sakamoto et al., 2005). This suggests a favorable therapeutic window for hBD-2 before unacceptable toxicity becomes an issue. Clearly, next steps in conclusively showing the efficacy of hBD-2 against CoV-2 would be to conduct live viral *in vitro* infections of ACE2 expressing cells in a BSL3 facility followed by *in vivo* CoV-2 infection studies in appropriate animal models (Kim et al., 2020).

*In vivo* application of hBD-2 has proven successful in addressing a number of diseases. This includes a recent study demonstrating efficacy in experimental colitis in a mouse model (Koeninger et al., 2020) and therapeutic intranasal application of hBD-2 to reduce the influx of inflammatory cells into bronchoalveolar lavage fluid (Pinkerton et al., 2020). Of relevance to our study is the use of smaller hBD fragments; i.e., mimetics, of mouse beta defensin 4 (mBD-4) (Zhao et al., 2016), the ortholog of hBD-2, that when administered intra-nasally, rescued 100% of mice from the lethal challenge of human and avian influenza A, SARS-CoV and MERS-CoV (LeMessurier et al., 2016). Therefore, should *in vivo* studies of hBD-2 prove to be successful in blocking live CoV-2 infection in an animal model, the fact that the peptide is endogenous to humans and would not elicit an immunogenic response, give it a high probability of being safe and a quicker route to human clinical trials. In fact, several AMPs, as well as AMP mimetics are currently undergoing clinical trials for multiple different diseases (Mookherjee et al., 2020).

A recent *in silico* molecular docking study predicted a strong binding interaction between LL-37 and the RBD, demonstrating the blocking potential of LL-37 for ACE2 binding (Lokhande, 2020). This was followed up by a surface plasmon resonance study confirming the simulation results (Roth et al., 2020). Since LL-37 has also been shown to possess antiviral activity (Tripathi et al., 2015), these results support the idea that more than one AMP could be utilized, possibly in a “cocktail” to act as a potent viral blocking agent. Recent findings also highlight that neuropilin-1 (NRP1), a receptor involved in multiple physiological processes and expressed on many cell types (Roy et al., 2017), is being utilized by CoV-2 to facilitate entry and infection (Cantuti-Castelvetri et al., 2020; Daly et al., 2020). Time will tell if blocking ACE2 alone will be enough to reduce CoV-2 infection and/or reduce the severity of symptoms, or if an additional strategy of also blocking entry via NRP1 will be required.

Not unexpectedly, CoV-2 is mutating, albeit at a relatively slower rate than influenza viruses; i.e., two to six fold slower over a given time frame (Manzanares-Meza and Medina-Contreras, 2020). Ongoing studies indicate that it has developed a number of mutations of which 89 have been associated with the RBD (Chen et al., 2020; Wang et al., 2020). Furthermore, 52 out of 89 mutations are in the receptor-binding motif (RBM), i.e., the region of RBD that is in direct contact with ACE2, indicating that the virus may be accumulating mutations in that region to improve its interaction with ACE2 (Li et al., 2020). Fortunately, while these and other mutations appear to have evolved for greater transmissibility, they have not resulted in greater pathogenicity. The variant that has recently received much attention is “VUI-202012/01,” the one first reported in southeast England that presents with multiple amino acid changes to the spike protein (https://virological.org/t/preliminary-genomic-characterisation-of-an-emergent-sars-cov-2-lineage-in-the-uk-defined-by-a-novel-set-of-spike-mutations/563) While not confirmed yet in animal experiments, early reports suggest that it may be >50% more infectious than the parent strain. Of particular importance to us is the asparagine to tyrosine conversion in position 501 (N501Y), as this is one of the contact residues within the RBM that plays a role in binding to ACE2. As shown in Figure S7, the bound hBD-2 monomer and dimer are on average not close to the sidechain site of residue 501 (> 8A between nearest atoms). Furthermore, compared to the ring-ring (pi-pi) contact between residue side-chains, which is highly probable between the UK-mutant RBD and ACE2, stabilizing the interaction as shown by a deep mutagenesis study with the N501Y mutation enhancing binding (Starr et al., 2020), neither a sidechain ring or positively charged sidechain of hBD-2 appears to come near in our models of its complex with the (original) RBD. At the same time it should be noted that the interaction of the RBD with ACE2, and especially with hBD-2, is considerably dynamic (Zhang et al., 2016; Zhang and Buck, 2017). Although this has not yet been measured in the RBD:ACE2 or RBD:hBD-2 platforms, the entropy of the interaction is likely to be not as unfavorable as seen in complexes where one or both partner proteins have to become significantly rigid. It is now becoming clear that many protein-protein complexes are inherently dynamic (Zhang et al., 2016; Zhang and Buck, 2017), thus minimizing the unfavorable entropy change that would otherwise occur on binding. This is especially important for the binding of peptides, which may be relatively unstructured in solution and suggests that design of hBD-2 and LL-37 derived peptides would be a fruitful endeavor.

While vaccines against SARS-CoV-2 have recently been approved by the FDA and are planned for distribution and administration in a large scale to cover most of the American population over the next year, we see the AMP strategy as complementary to vaccines. While the CoV-2 vaccines appear to show >90% efficacy, there will certainly be some degree of morbidity and mortality, as seen in all vaccines (Kaselitz et al., 2019), many people will refuse vaccination (Pogue et al., 2020; Schwarzinger et al., 2010) and a significant number will either fail to mount effective neutralizing antibodies or high enough titers (Goodwin et al., 2006; Ndifon et al., 2009; Ovsyannikova et al., 2017). Many of these low or non-responders are predicted to be in the COVID-19 high-risk population. Additionally, vaccines more than likely will provide protection for a limited amount of time, as neutralizing antibodies wane, and many people could face reinfection. Because of the multiple advantages of using small peptides like hBD-2 and their derived smaller mimetics, such as high specificity, low toxicity, lack of immunogenicity, low cost of production and ease of administration, they possess the potential for both safety and efficacy. Molecules such as hBD-2 could be delivered, in the future, intra-orally and/or intra-nasally as prophylactic aerosols, in early stages of infection, when telltale symptoms appear and in combinatorial therapeutic approaches for more severe situations.

## ACKNOWLEDGEMENTS

We thank Energy Center (CESR) of Tennessee Technological University for partially supporting graduate student Jackson Penfield and the pilot fund from Drs. Weinberg and Buck for undergraduate student Ann Brewer. The simulations were mainly done on Ohio Supercomputer Center Pitzer machines, and partly on high performance computers in Tennessee Technological University. We thank Dr. Jesse Bloom, Fred Hutchinson Cancer Center for kindly proving the plasmids to generate Spike pseudovirus and HEK 293T cells expressing ACE2 receptor, and Dr. Parvesh Shrestha of the Buck lab, for help with MST experiments. Dr. Buck is currently funded by NIH R01 grant from the National Eye Institute R01EY029169 and his part of the project was also supported by pilot grant from the Department of Physiology and Biophysics of Case Western Reserve University. Dr. Ramakrishnan is supported by NIH/NIAID grants R01AI116730 and R21AI144264, NIH/NCI grant R21 CA246194 and a pilot funding from NORD Family Foundation for COVID related research. Dr. Weinberg was supported by pilot funds from the Department of Biological Sciences of the School of Dental Medicine, CWRU.

## AUTHOR CONTRIBUTIONS

Conceptualization, LZ, SKG, PR, MB, AW. Methodology, LZ, SKG, PR, MB. Investigation, LZ, SCB, JM, JP, AB, SKG, PR. Writing – Original Draft, LZ, SKG, PR, AW. Writing – Review & Editing, SKG, PR, LZ, MB, AW. Visualization, SKG, MB, PR, AW. Supervision, LZ, PR, MB, AW, Project Administration, AW. Funding Acquisition, LZ, MB, PR, AW.

## METHODS

### Resource availability

Further information and requests for resources and reagents should be directed to and will be fulfilled by the Lead Contact, Aaron Weinberg.

### Materials Availability

This study did not generate new unique reagents.

### Cells

HEK 293T and HEK 293T cells stably expressing ACE2 receptor (ACE2 HEK293T) were cultured in DMEM media containing 10% FBS, 100 U/ml penicillin/streptomycin and 4 mM L-Glutamine.

### Plasmids

pHAGE-CMV-Luc2-IRES-ZSgreen-W, HDM-HgPM2, HDM-tat1b, pRC-CMV-Rev1b, and SARS-CoV-2 Spike-ALAYT plasmids were previously described (Crawford et al., 2020). FLAG and HIS tagged RBD were expressed from a pcDNA3 vector with leader sequence and leucine zipper as previously described (Ramakrishnan et al., 2004).

### Structure information

The structure of human beta defensin 2 (hBD-2) in the monomer and dimer form is available in the PDP with ID 1FD3 (Hoover et al., 2000). The hBD-2 sequence is 41 residues long: GIGDPVTCLKSGAICHPVFCP**RRYKQ**IGTCGLPGTKCCKKP. The five boldened residues were found to form hydrogen bonds with the RBD during the simulations (see below/main paper). The structure of the <u>RBD</u> domain of the Spike protein is also available in complex with ACE2 at 2.45 Å resolution in the PDB with ID 6M0J (Lan et al., 2020)

## METHOD DETAILS

### Docking and all-atom simulations

Two kinds of docking programs were applied; one was Cluspro (Kozakov et al., 2006; Kozakov et al., 2017; Porter et al., 2017; Vajda et al., 2017), while the other was HADDOCK (Dominguez et al., 2003; van Zundert et al., 2016). The x-ray structures of hBD-2 and of the SARS-CoV-2 S-protein RBD were uploaded to the Cluspro docking webserver without additional preparation. The best docked structures were clustered, with most of them showing that the hBD-2 binds to the RBD at sites used for the association between ACE2 and RBD. The best structure was selected based on the docking programs’ score and the predicted binding sites between hBD-2 and RBD. Cluspro is a rigid body protein docking method. It is based on a Fast Fourier Transform correlation approach, which makes it feasible to generate and evaluate billions of docked conformations by simple scoring functions as shown in Equation (1). It is an implementation of a multistage protocol: rigid body docking (used PIPER), an energy based filtering, ranking the retained structures based on clustering properties, and finally, the refinement of a limited number of structures by energy minimization. In the Cluspro docking, the PIPER interaction energy is calculated using the following equation:

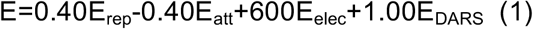

Here, E_rep_ and E_att_ are contributions of the van der Waals interaction energy, and E_elec_ is an electrostatic energy term. E_DARS_ is a pairwise structure-based potential constructed by the Decoys as the Reference State (DARS) method (Chuang et al., 2008). It primarily represents a desolvation contribution, i.e., the free energy change due to the removal of the water molecules from the interface (Kozakov et al., 2006). Since in the PIPER calculation, the entropic term was not included in Cluspro docking, the PIPER energy result should not be used to rank clusters. Instead, the population of clusters was applied to rank the clusters. In our simulations, the RBD:hBD-2 complex structure from the top cluster was taken and continued with all-atom molecular dynamics simulations.

In the HADDOCK docking, since the binding interface between the ACE2 receptor and RBD are known, residues from 400 to 520 on the RBD were selected as the target binding sites, while the entire hBD-2 peptide taken as a potential binding site. Default values for all other parameters were applied. After that, the best 5 structures, by HADDOCK scoring, were selected.

Based on the best 6 (including above 5 from HADDOCK and 1 from Cluspro docking) structures predicted above, all-atom molecular dynamics simulations were set up using the CHARMM36m (Huang et al., 2017) forcefield and VMD program (Humphrey et al., 1996). One of the deprotonated states of histidine was used (denoted HSD), and the native disulfide bonding in the hBD-2 was set up. After solvating the protein with an equilibrated box of TIP3P water molecules, the closest distance between atoms on the proteins and the edge of simulation box is 12 Å. The equivalent of 0.15 M in Na and Cl ions was added into the box plus several ions to neutralize the net charge of the system. The desired temperature is 310 K and pressure is 1 atm, using standard thermo- and barostats. After a brief energy minimization using the conjugate gradient and line search algorithm, 4 ps of dynamics was run at 50 K, and then the system was brought up to 310 K over an equilibration period of 1 ns using NAMD program version 2.12 (Phillips et al., 2005). This was followed by trajectories that continued for up to 200 or 500 ns at 1 atm and 310 K using the NPT ensemble.

As a comparison, we also simulated the RBD bound with ACE2 using the structures from (Lan et al., 2020) and the same method as above. HBD-2 can also form a non-covalent dimer at high concentration in solution (Hoover et al., 2000) (with PDB ID of 1FD3). The initial bound structure of the hBD-2 dimer with the RBD was predicted using targeted HADDOCK docking. The best structure predicted was used in all-atom MD simulations as detailed above.

The simulation systems, set up, the number of atoms and box size information are shown in Table S1.

To analyze the trajectories, the Root Mean Square Deviation (RMSD) and Fluctuations (RMSF) of the proteins were calculated using the VMD program and an in-house analysis script based on the coordinates of the backbone Ca atoms after aligning the trajectories respectively, to the original crystal structure of the RBD, hBD-2, and to the initial complex structure of the RBD and hBD-2 predicted from docking. The buried surface area (BSA) for the complex was calculated in two steps using the VMD program and a script using the Richards and Lee method with the water probe size of 1.4 Å (Lee and Richards, 1971). First, the total solvent accessible surface area of the complex (ASA_complex_) was calculated based on the complex’s trajectory. Second, the accessible surface area of each protein in the complex (ASA_rbd_, ASA_hbd2_) was calculated for each protein individually. Then, the BSA is calculated using Equation (2):

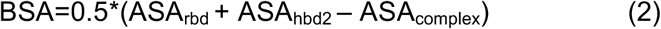

The number of hydrogen bonds between the RBD and ACE2 or the RBD and hBD-2 were calculated using the VMD program with the heavy atom distance cutoff of 3.0 Å and the angle cutoff of 20 degrees deviation from H-bond linearity. The time a particular H-bond is formed over the course of the simulation is monitored and is expressed as % occupancy. In order to find out the residues on the binding interface, the closest distance between every residue atom (including hydrogen) between the RBD and hBD-2 was calculated and averaged over the trajectory run. The average distances between each residue on RBD and on hBD-2 are shaded by proximity on a red to white color-scale and were used to build the distance maps. Furthermore, based on the long term simulation trajectories of the complexes of Supplementary Table S1, the total pairwise interaction energy was calculated using the MM-GBSA method (Genheden and Ryde, 2015) by applying NAMD and the NAMD energy plugin of the VMD program(Humphrey et al., 1996). This interaction energy (E_binding) is calculated using Equation (3):

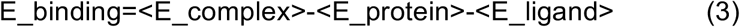

E_complex is the potential energy of protein-ligand complex, E_protein is the potential energy of protein, and E_ligand is the potential energy of ligand. < > is the ensemble average over simulation time. In the MM-GBSA method, the solvent effect was counted using the generalized Born implicit solvent model (GBIS)(Tanner et al., 2011).

### Measurement RBD:ACE2 association *in vitro*

Untagged hBD-2 and N-terminally His-tagged RBD were purchased from Peprotech, Inc. and Raybiotech Inc., respectively. Binding experiments were carried out with a Monolith NT.115 Microscale Thermophoresis (MST) instrument (NanoTemper, Inc.) at room temperature in pH 7.1 phosphate buffer saline with 0.1% Tween-20 (PBS-T 0.1%). The RBD was labeled using the NanoTemper Monolith HIS-Tag Labeling Kit RED-trisNTA which labels His-tags with a fluorescent group. 40 nM of this RBD was mixed with a serial dilution of unlabeled hBD-2 in 0.2 mL micro reaction tubes (NanoTemper, Inc.) and then transferred to premium capillaries (NanoTemper, Inc.). The experiment was done with a triplicate set of tubes. Microscale thermophoresis monitors the change of the diffusion of proteins/peptides in microscopic temperature gradients upon protein binding. The dissociation constant K_d_ was obtained by fitting the binding curve with the quadratic solution for the fraction of fluorescent molecules that formed the complex between proteins A and T, calculated from the law of mass action K_D_ = [A]*[T]/[AT] where [A] is the concentration of free fluorescent molecule and [T] the concentration of free titrant and [AT] the concentration of complex of A and T. We also carried out the experiment with a labeled RBD as well as hBD-2 sample which had its disulphide bonds reduced by addition of 2.0 mM DTT, showing that disulphide bonds are essential for maintaining the folded structures (Hati and Bhattacharyya, 2020) and that these are required for the reasonably strong protein-protein interactions (Figure 5A).

### ELISA based assay

100 **μ**l of rhBD-2 (Peprotech, Inc.) (concentration as indicated in Figure 5B) in assay diluent buffer 2 (R&D system), were incubated in an RBD coated plate (Ray biotech, Inc.) at 4^0^C for 18 hrs. Plates were then washed 4 times with 300 **μ**l of wash buffer (R&D Systems, Inc.) followed by incubation with 100 **μ**l of biotinylated anti hBD-2 (Peprotech, Inc.)[0.1**μ**g/ml] for 1 hr. Plates were then washed again as stated above, incubated with 100**μ**l of Streptavidin-HRP (R&D system, Inc.) for 20 minutes. Signal was developed using TMB substrate and measured at 450nm using a microplate reader.

### Immunoprecipitation and Western blotting

To study interaction between hBD-2 and His-RBD, recombinant hBD-2 (Peprotech, Inc.) with or without recombinant HIS-tagged-RBD (Sino Biologicals, Inc.) were pre-incubated at room temperature for 1 h in binding buffer (30mM HEPES pH 7.6, 5mM MgCl2, 150mM NaCl, 0.5mM dithiothreitol, 1 % Triton X-100 and 1mM EDTA) and then incubated with washed Ni-NTA agarose resin beads (25 µl) overnight at 4°C. Beads were collected by centrifugation at 1000 rpm for 1 min and washed thrice with binding buffer. Beads were boiled with 30 µl of Laemmli sample buffer and were analyzed by Western blotting (WB). Briefly, samples were separated on 20% SDS-Polyacrylamide gels and proteins were then transferred to nitrocellulose membrane (0.2 µm pore size) at 70V for 40 min in cold. Membranes were blocked with 5% milk in TBST and then probed with goat anti-human BD2 antibody (0.2 µg/ml; Peprotech), followed by secondary antibody (1:5000) at room temperature, and visualized by enhanced chemiluminescence. To study the ability of hBD2 to compete with RBD binding to ACE2, ACE2 HEK 293T cells were seeded in 6 cm plates. At 50% confluency, media was replaced with conditioned media from HEK 293T cells transfected with secreted FLAG RBD plasmid or control media in the presence or absence of hBD2 (1.0 and 3.0 µg/ml) and incubated at 37°C for 30 min. Cells were washed and collected in PBS-EDTA solution and then lysed in Triton lysis buffer. Lysates were centrifuged at 12000 g for 10 min at 4°C, and immunoprecipitated using M2 FLAG beads (Sigma) for 2 hours at 4°C. Beads were collected, washed, and boiled with Laemmli sample buffer and analyzed by Western blotting.

### CoV-2 spike-pseudotyped luciferase assay

Pseudotyped SARS-CoV-2 spike virus was generation and luciferase assay was carried out as described previously (Crawford et al., 2020). Briefly, HEK 293T cells were transfected with luciferase-IRES-ZSgreen, HDM-HgPM2, HDM-tat1b, PRC-CMV-Rev1b, and SARS-CoV-2 Spike-ALAYT plasmids as described (Crawford et al., 2020) Culture supernatants were harvested 48 hours after transfection and used to infect ACE2 HEK293T cells. To study the effect of hBD2 on spike pseudotyped virus entry, ACE2 HEK 293T cells were incubated with pseudovirions and varying concentration of HBD2 (0-15 µg/ml) for 48 hours. Cells were lysed and luminescence was measured using luciferase assay system following manufacturer’s instructions (Promega, Inc.) in Spectramax i3 microplate detection platform (Molecular Devices, Inc.).

