## Supplementary information for "Molecular dynamics simulations and functional studies reveal that hBD-2 binds SARS-CoV-2 spike RBD and blocks viral entry into ACE2 expressing cells"

**Table S1:** Simulation name, initial structure source, number of atoms, box size and simulation length of simulations performed in this project.

| Simulation name | Initial structure source | Number of atoms | Box size (Å$^3$) | Simulation length (ns) |
|---|---|---|---|---|
| RBD:ACE2 | (Lan et al., 2020) | 104857 | 85.1x95.2x138.9 | 50 |
| RBD:hBD-2/c | Cluspro docking | 40524 | 75.0x72.8x79.8 | 500 |
| RBD:hBD-2/h1 | Haddock docking | 32849 | 88.5x69.4x57.9 | 200 |
| RBD:hBD-2/h2 | Haddock docking | 32468 | 94.7x68.1x54.6 | 500 |
| RBD:hBD-2/h3 | Haddock docking | 38330 | 87.0x72.9x65.1 | 200 |
| RBD:hBD-2/h4 | Haddock docking | 35126 | 93.4x71.9x56.4 | 200 |
| RBD:hBD-2/h5 | Haddock docking | 39179 | 87.7x71.4x67.4 | 100 |
| RBD:hBD-2/c1 | RBD:hBD-2/c final | 32468 | 94.7x 68.1x49.5 | 500 |
| RBD:hBD-2/c2 | RBD:hBD-2/c final | 32468 | 94.7x68.1x49.3 | 500 |
| RBD:hBD-2/c3 | RBD:hBD-2/c final | 32468 | 94.7x68.1x49.3 | 200 |
| RBD:hBD-2-dimer | Haddock docking | 45913 | 74.3x72.8x91.3 | 500 |
| ACE2 in solvent | (Lan et al., 2020) | 70740 | 93.3x94.8x78.6 | 50 |
| RBD in solvent | (Lan et al., 2020) | 36044 | 65.7x73.3x74.3 | 500 |
| hBD-2 in solvent | (Hoover et al., 2000) | 11511 | 53.5x52.5x40.7 | 500 |
| hBD-2 dimer in solvent | (Hoover et al., 2000) | 15455 | 58.9x47.9x54.2 | 500 |

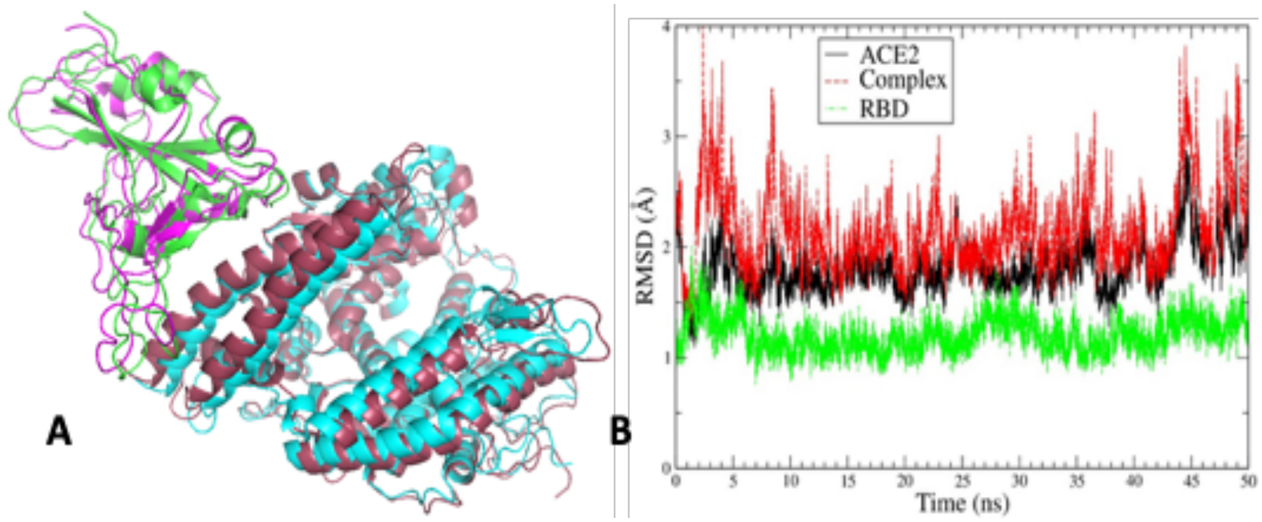


**Figure S1. Reference for molecular dynamics (MD) simulations of RBD:ACE2 – supportive information for Fig. 1.** [A] Comparison of the initial structures (shown in cyan for ACE2 and green for RBD) and last structure (shown in raspberry for ACE2 and magenta for RBD) after 50 ns all-atom MD simulation for the RBD from SARS-COV-2 spike protein in complex with ACE2. [B] Ca RMSD for RBD, ACE2 and RBD:ACE2 complex as a function of simulation time

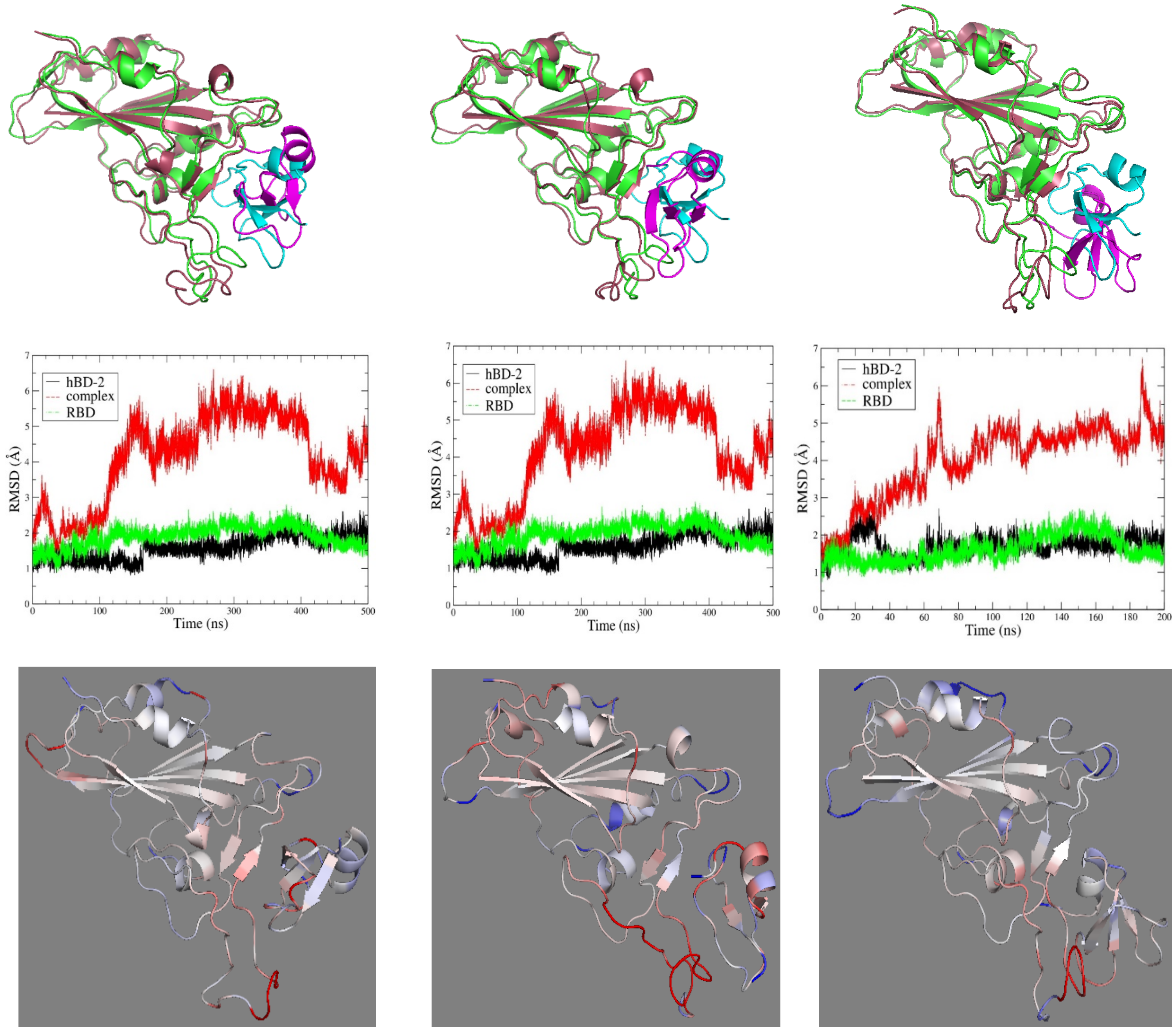


RBD:hBD-2/c1 RBD:hBD-2/c2 RBD:hBD-2/c3

**Figure S2. Additional MD simulations of the RBD : hBD-2 complex.** RBD:hBD-2/c1, /c2 and /c3 show similar trends to simulation RBD:hBD-2/c (Fig. 2). **Top:** Final structures after all-atom simulation started with different seeds for velocity assignment, superimposed on initial structure. **Middle:** Color code as Fig. 2. RMSD of proteins in the complex and of the complex itself. **Bottom:** Difference in RMSF of proteins in complex to their fluctuations when unbound.

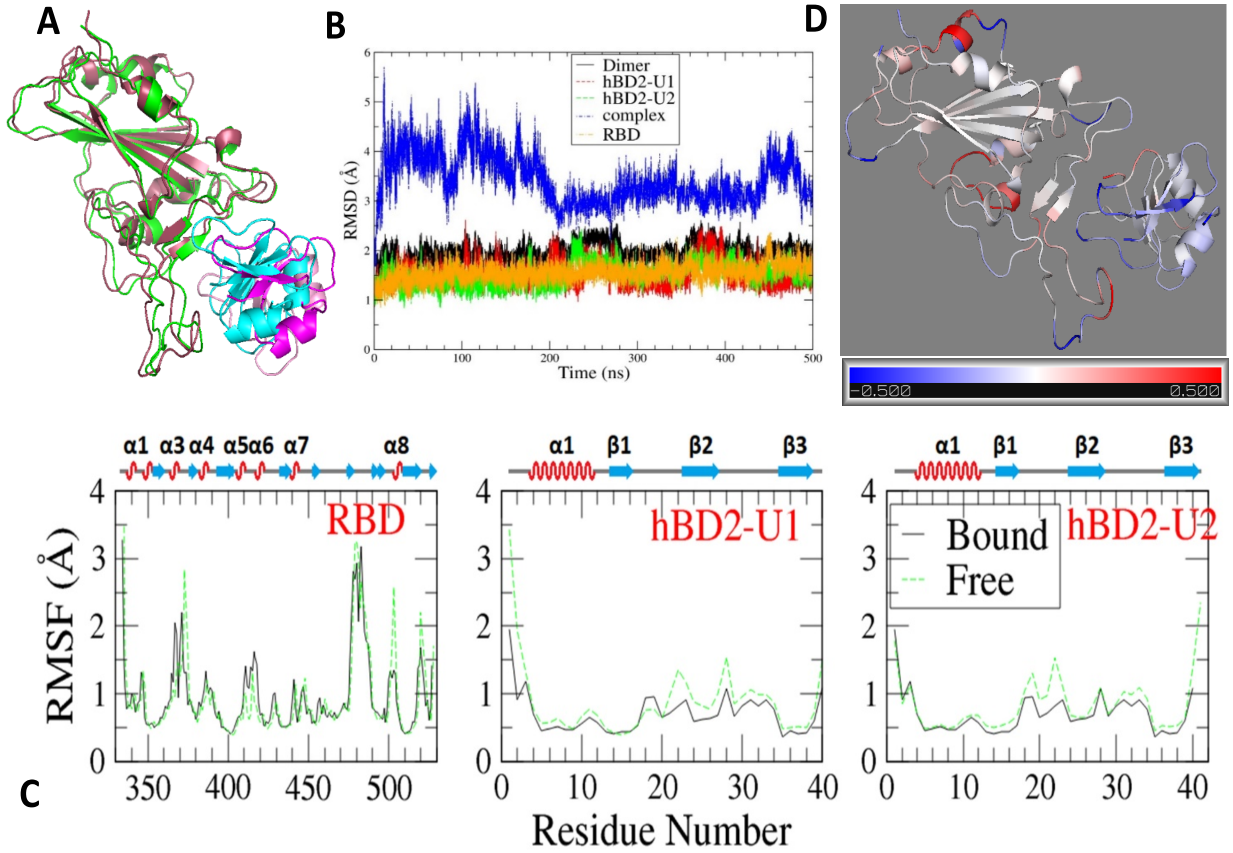


**Figure S3**. **MD simulations of the RBD:hBD-2 complex show that the hBD-2 dimer also forms a stable complex with the RBD.[A]** Comparison of the initial and last structure over 500 ns simulation (shown in cyan for hBD-2 dimer and green for RBD and in magenta for hBD-2 dimer and raspberry for RBD, respectively). **[B]** RMSD of the individual proteins in the complex and of the complex itself. **[C]** RMSF of RBD (left) and hBD-2 dimer unit 1 (middle) and unit 2 (right) in the complex over 500 ns in comparison with values for the unbound (free) proteins. The secondary structure of ACE2 and RBD are indicated. **[D]** Difference in RMSF between bound and free proteins. The data are mapped to the cartoon representation of the complex with color bar (Bottom) indicating the range of -0.5 Å (in blue) to 0.5 Å (in red)

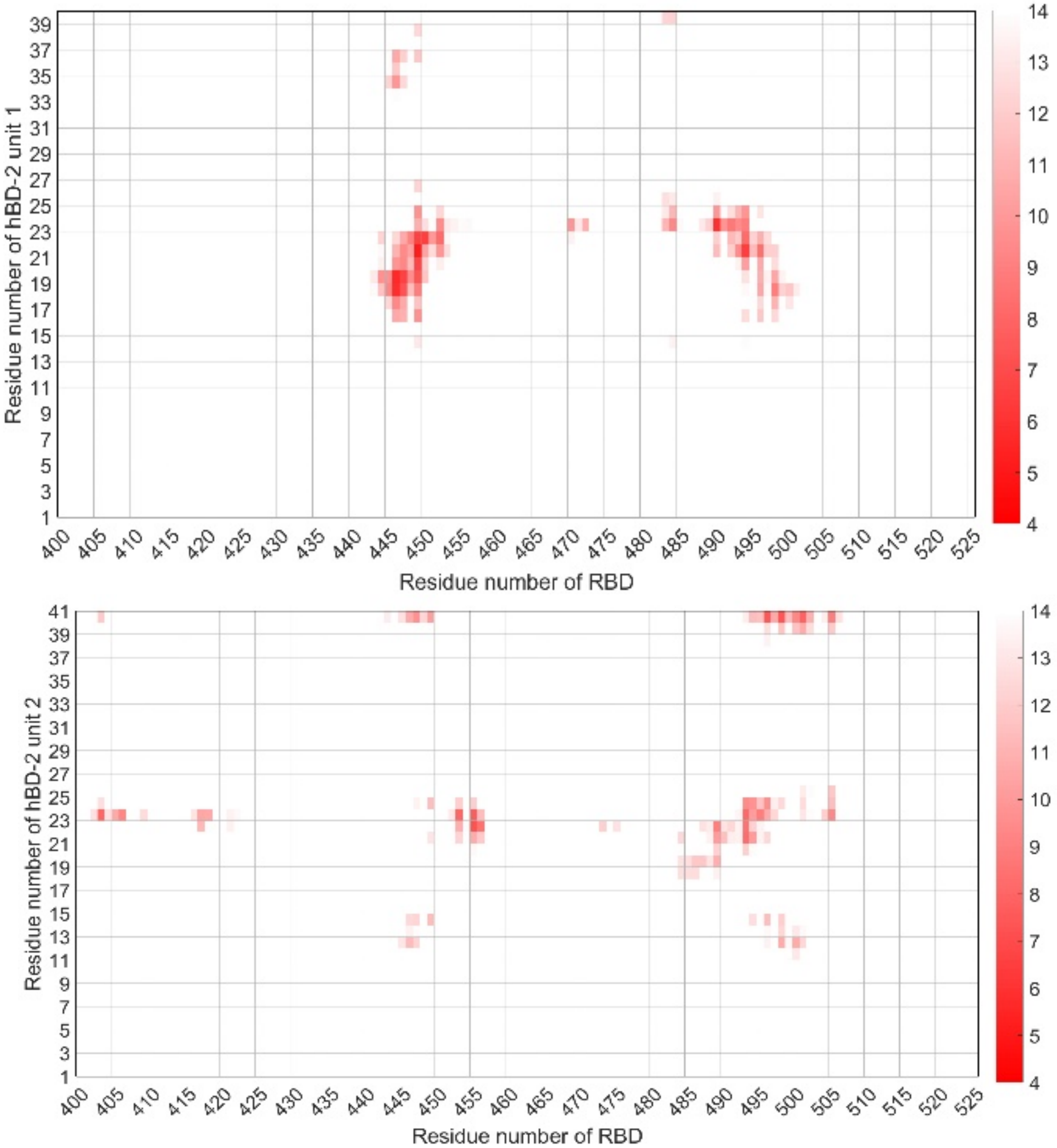


**Figure S4. Distance map of average inter-protein residue proximity between the first unit (Top) and the second unit (Bottom) of hBD-2 dimer with RBD over 500 ns simulation.** The residue pair distances are color coded by average proximity over the length of the simulations (see color scale, right).

**A**

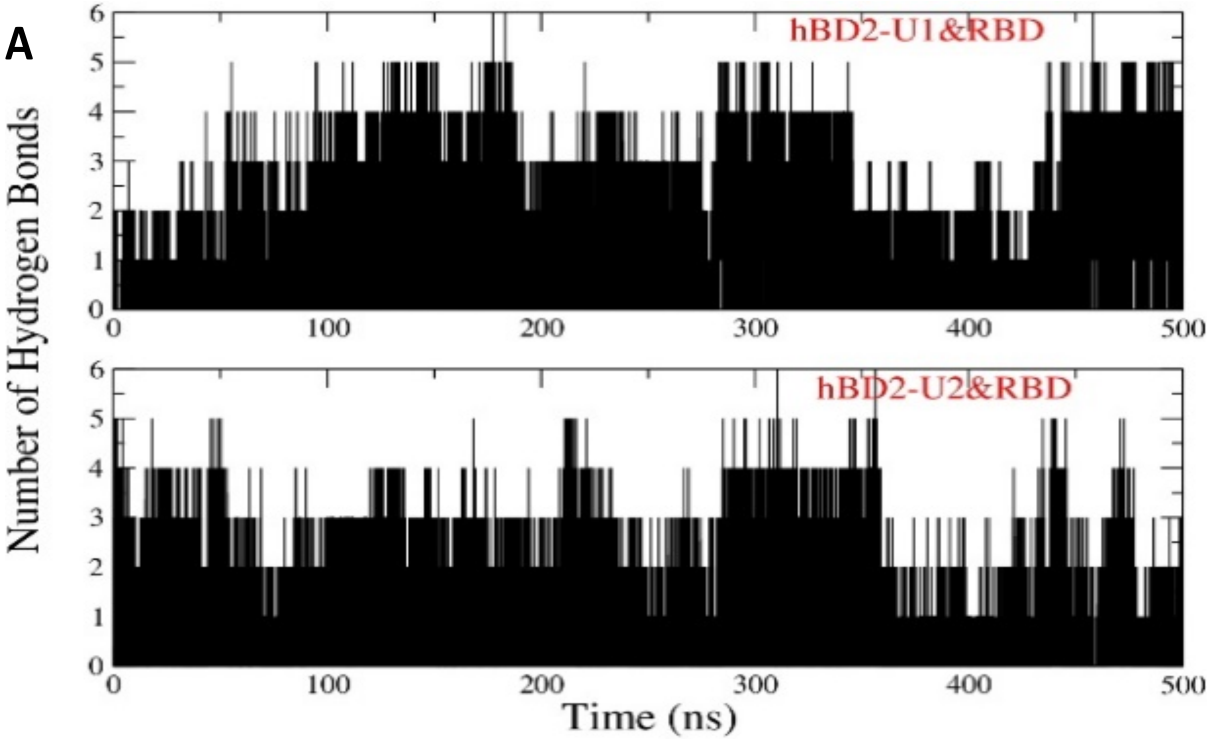


**B**

| Donor | Acceptor | Occupancy |
|---|---|---|
| LYS40-Side (hBD-2 U1) | GLU484-Side (RBD) | 43.81% |
| ARG23-Side (hBD-2 U1) | GLU484-Side (RBD) | 47.98% |
| TYR449-Side (RBD) | PHE19-Main (hBD-2 U1) | 11.24% |
| ARG23-Side (hBD-2 U2) | GLU406-Side (RBD) | 62.05% |

**Figure S5. Hydrogen bonds between RBD and dimeric hBD-2 are more stable compared to those with monomeric hBD-2 (compare with Fig. 3).**
**[A]** Number of hydrogen bonds formed between each unit of the hBD-2 dimer with the RBD. **[B]** Details of residues on RBD and hBD-2 units as donor and acceptor forming hydrogen bonds.

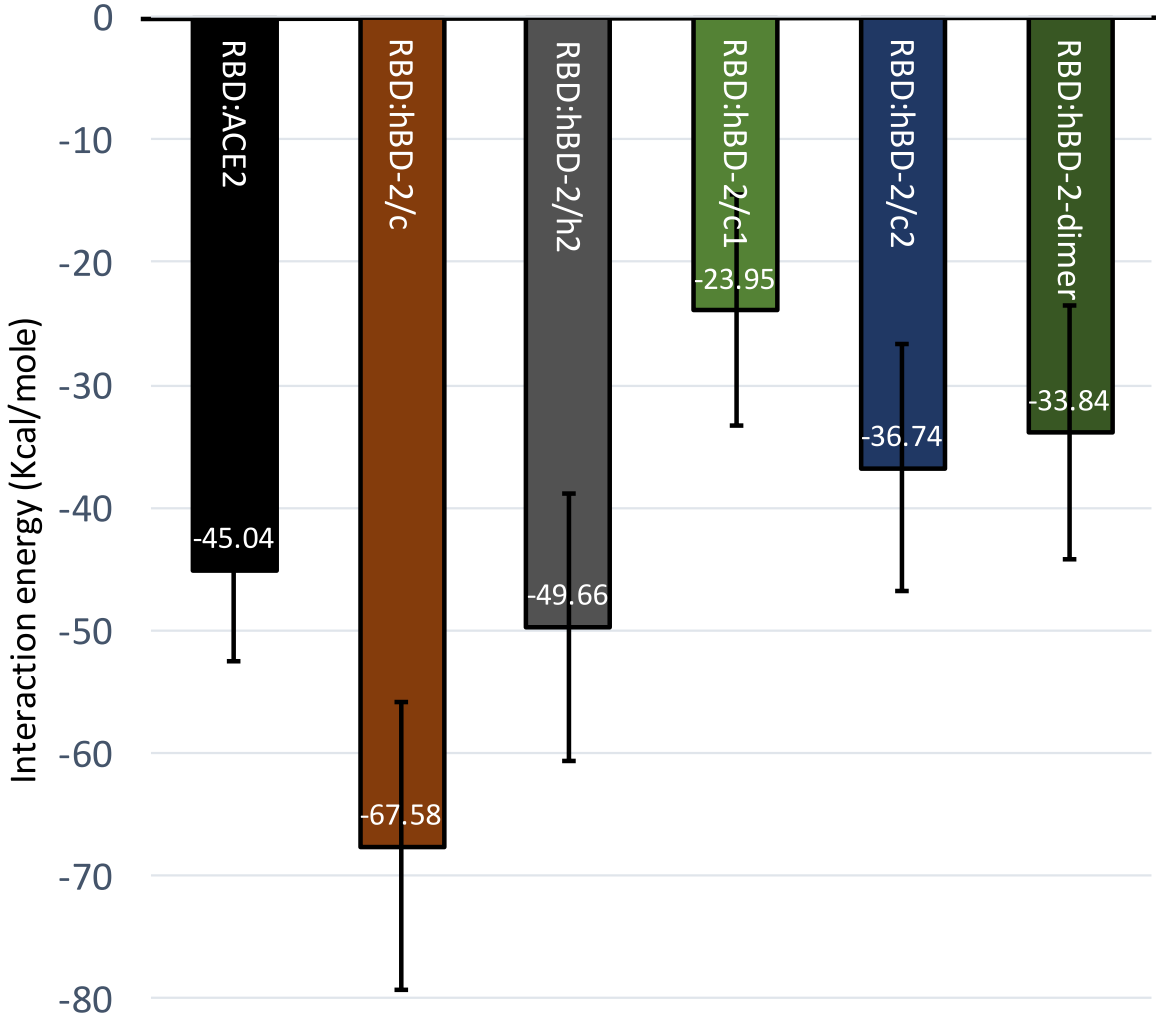


**Figure S6. Histogram representation of estimated binding energies of hBD-2 binding to the RBD vs. RBD binding to ACE2.**

Comparison of interaction energies for ACE2, hBD-2 monomer/dimer binding with RBD. E_binding = E_complex - E_protein - E_ligand (see Methods for details)

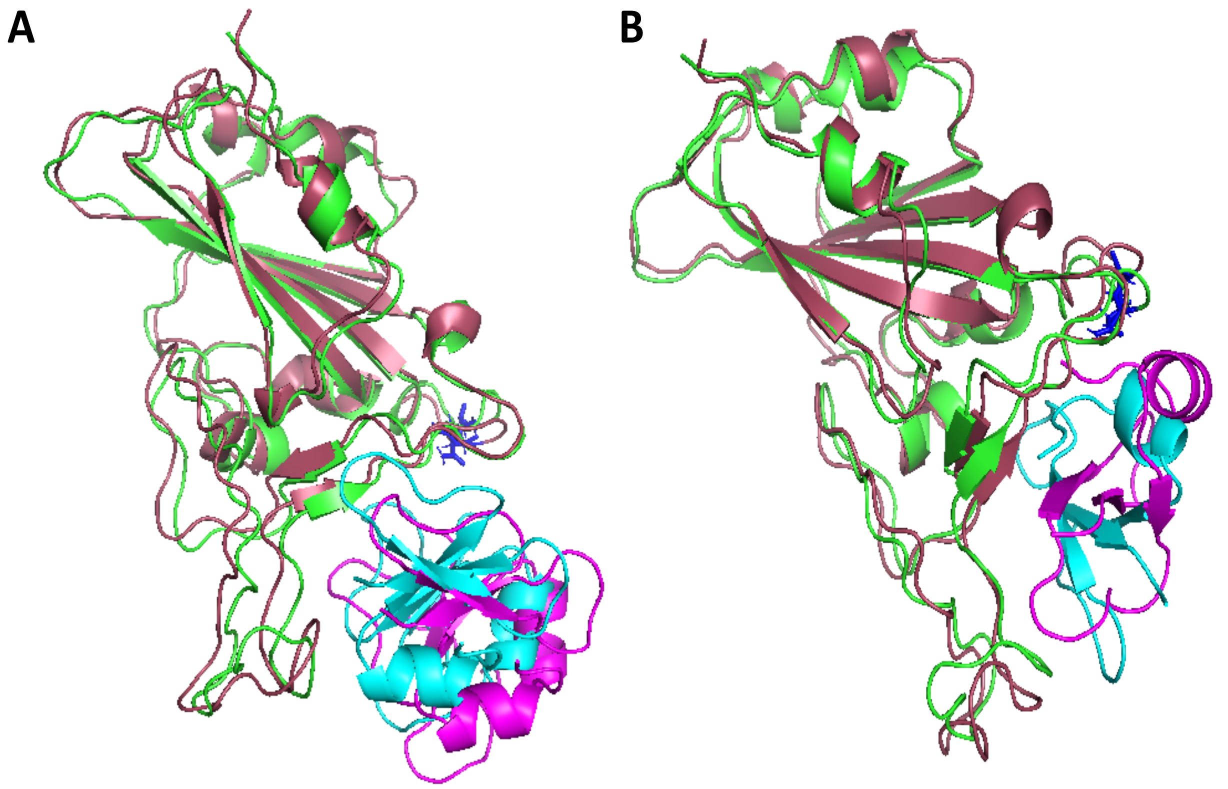


**Figure S7: Location of the N501Y mutation in RBD relative to bound hBD-2 monomer does not suggest additional interactions.** RBD binding with hBD-2 dimer (simulation RBD:hBD-2-dimer) [**A**] and with hBD-2 monomer (simulation RBD:hBD-2/h2) [**B**] by superposition of initial and final structures (color code as in Fig. S1). The N501 residue highlighted in blue sticks. Both structures show that the N501 residue is at the corner of the binding sites of hBD-2 with RBD; i.e., not within the main region of interaction.
